# Phosphoregulation of DSB-1 mediates control of meiotic double-strand break activity

**DOI:** 10.1101/2022.02.16.480793

**Authors:** Heyun Guo, Ericca L. Stamper, Aya Sato-Carlton, Masa A. Shimazoe, Xuan Li, Liangyu Zhang, Lewis Stevens, KC Jacky Tam, Abby F. Dernburg, Peter M. Carlton

## Abstract

In the first meiotic cell division, proper segregation of chromosomes in most organisms depends on chiasmata, exchanges of continuity between homologous chromosomes that originate from the repair of programmed double-strand breaks (DSBs) catalyzed by the Spo11 endonuclease. Since DSBs can lead to irreparable damage in germ cells, while chromosomes lacking DSBs also lack chiasmata, the number of DSBs must be carefully regulated, so as to be neither too high nor too low. Here, we show that in *Caenorhabditis elegans*, meiotic DSB levels are controlled by the phosphoregulation of DSB-1, a homolog of the yeast Spo11 cofactor Rec114, by the opposing activities of PP4^PPH-4.1^ phosphatase and ATR^ATL-1^ kinase. Increased DSB-1 phosphorylation in *pph-4*.*1* mutants correlates with reduction in DSB formation, while prevention of DSB-1 phosphorylation drastically increases the number of meiotic DSBs both in *pph-4*.*1* mutants as well as in the wild type background. *C. elegans* and its close relatives also possess a diverged paralog of DSB-1, called DSB-2, and loss of *dsb-2* is known to reduce DSB formation in oocytes with increasing age. We show that the proportion of the phosphorylated, and thus inactivated, form of DSB-1 increases with age and also upon loss of DSB-2, while non-phosphorylatable DSB-1 rescues the age-dependent decrease in DSBs in *dsb-2* mutants. These results suggest that DSB-2 evolved in part to compensate for the inactivation of DSB-1 through phosphorylation, to maintain levels of DSBs in older animals. Our work shows that PP4^PPH-4.1^, ATR^ATL-1^, and DSB-2 act in concert with DSB-1 to promote optimal DSB levels throughout the reproductive lifespan.

## Introduction

To reduce chromosome number from diploid to haploid during sexual reproduction, homologous chromosomes must segregate to different daughter cells in the first division of meiosis. Most organisms achieve this segregation by linking homologous chromosomes with chiasmata, exchanges of continuity between chromatids that derive from repair of programmed double-strand breaks (DSBs). DSBs are created by the conserved endonuclease Spo11 acting in concert with an array of cofactors (1–4). The initiation of DSBs needs to be strictly controlled, due to their deleterious potential: not only can unrepaired breaks lead to apoptosis (5, 6), but unfavorable repair mechanisms such as non-homologous end-joining or non-allelic homologous recombination acting on DSBs (7) can lead to genome rearrangement or deletions. Despite these dangers, however, every chromosome pair requires at least one crossover for proper segregation, so DSB initiation must be allowed to occur until this condition has been met. Accordingly, DSBs must be regulated in space and time to achieve a number that is not too high, but not too low. How this regulation occurs, i.e., how each species enforces the correct level of DSBs they require (8), remains an unsolved mystery.

A large body of work has shown that the DNA damage sensor kinases ATM and/or ATR control DSB initiation and repair at multiple levels in mammals (9), Drosophila (10), budding yeast (11–13) and other organisms. When a break occurs, ATM/ATR (yeast Tel1/Mec1) locally phosphorylate many substrates, including the Spo11 accessory protein Rec114, leading to a local reduction in further DSB formation in budding yeast (11), a phenomenon known as DSB interference (12, 14). ATR(Mec1) activity induced by replication stress has also been shown to reduce the chromosome loading of Rec114 in budding yeast (15). In mice, both ATM and ATR kinase act to remove recombination factors from the vicinity of DNA breaks and suppress DSB initiation (9, 16). However, in both cases it is unknown whether any phosphatase counteracts or regulates the anti-DSB activity of these kinases.

Rec114, originally discovered through screens in budding yeast to identify genes required for initiation of meiotic recombination (17, 18), acts in concert with Mei4 and Mer2, together referred to as the RMM complex, to promote DSB initiation (19, 20). Homologs of yeast Rec114 include mouse Rec114 (20, 21), fission yeast Rec7 (22), and *C. elegans* DSB-1 and DSB-2, all of which are required for meiotic DSB formation (23, 24). A homolog of Mei4, DSB-3 in *C. elegans*, is also required for DSB formation (25), but no nematode homolog of Mer2 has been identified as of this writing. While the exact mechanism of DSB promotion by the RMM complex remains obscure, recent evidence suggests it may act as a scaffold for the Spo11 core complex (26).

In budding yeast and many other organisms, mutations that abolish DSB formation or processing also block synapsis, the polymerization of a protein macroassembly called the synaptonemal complex (SC) that holds chromosomes together in meiosis. This dependence has led to the conclusion that synapsis between homologous chromosomes is dependent on successful meiotic recombination in these organisms (27–31). In contrast, in *C. elegans* and *Drosophila melanogaster*, homologous chromosomes can pair and synapse in the complete absence of recombination (2, 32), and thus it has been suggested that these organisms achieve recombination-independent pairing and synapsis. However, while *C. elegans* can achieve homologous synapsis in the absence of DSB formation, recent evidence suggests this is equivalent to an early form of dynamic and unstable synapsis, which is normally later stabilized by DSB-induced recombination (33–35). The extent to which recombination contributes to homologous pairing and synapsis in *C. elegans* is not well understood.

Protein substrates phosphorylated by ATM and ATR are known to be dephosphorylated in many contexts by the highly-conserved serine/threonine Protein Phosphatase 4 (PP4) (36–39). Previously, we have shown that PP4 in *C. elegans* (PPH-4.1) is required for viability-supporting levels of DSB initiation as well as homologous pairing and synapsis during meiotic prophase (40).

In this work, we show that the DSB-promoting activity of DSB-1 is controlled by both PPH-4.1 and ATR (*C. elegans* ATL-1), and that meiotic DSB levels are decreased by the phosphorylation of DSB-1. During meiotic prophase, DSB-1 is phosphorylated in an ATL-1-dependent manner to inhibit DSB formation and protect the genome against excessive DSBs. In contrast, DSB-1 is dephosphorylated in a PPH-4.1-dependent manner, thereby promoting a number of DSBs sufficient to form a crossover on each chromosome pair for proper chromosome segregation. Since ATM/ATR kinases are known to be activated by DNA breaks, our model predicts that activated ATR kinase turns off DSB machinery via DSB-1 phosphorylation once sufficient levels of recombination intermediates are generated. This feedback mechanism could tune DSB levels while ensuring the formation of crossovers via phosphoregulation of DSB-1. Moreover, we find that the homologous pairing and synapsis defects in *pph-4*.*1* mutants are significantly rescued when DSB levels are increased by a non-phosphorylatable allele of *dsb-1*, adding to the growing evidence that DSBs can strengthen homologous synapsis in *C. elegans*. Our results shed light on fundamental mechanisms of meiotic chromosome dynamics regulated by contrasting kinase and phosphatase activities.

## Results

### DSB-1 undergoes phosphorylation which is prevented by PPH-4.1^PP4^ phosphatase

We previously showed that the activity of the PP4 phosphatase catalytic subunit, PPH-4.1, is necessary for normal levels of DSB initiation in *C. elegans* (40). This result suggested a hyperphosphorylated substrate acts to inhibit DSBs in *pph-4*.*1* mutants. Previous work in yeast showed that the Spo11 cofactor Rec114 is phosphorylated by ATM^Tel1^ and ATR^Mec1^ (11) in response to meiotic DSBs, and this phosphorylation suppresses DSB production.

In *C. elegans*, two recently-diverged orthologs of Rec114, DSB-1 and DSB-2, are also required for normal DSB formation (23, 24). To examine if PPH-4.1 regulates DSB levels through DSB-1, we determined whether loss of *pph-4*.*1* leads to DSB-1 hyperphosphorylation. We performed western blotting on extracts from animals carrying a GFP fusion of DSB-1 at the endogenous locus, comparing animals treated with RNAi against *pph-4*.*1* to control animals treated with an empty RNAi vector. The effectiveness of *pph-4* RNAi was verified by observing univalents in diakinesis oocytes (**Supplemental Figure 1A**). Two major bands were apparent in both treatments: one at the predicted size for DSB-1, and one more slowly-migrating (**Figure 1A**). In extracts from the *pph-4*.*1* RNAi-treated animals, the upper band was more intense, and the band at the expected size less intense. This result suggests that depletion of PPH-4.1 leads to hyperphosphorylation of DSB-1. We also probed blots from control and *pph-4*.*1* RNAi-treated worms with polyclonal antibodies directed against DSB-1 itself, and obtained similar results (**Supplemental Figure 1B**). To test whether the slow-migrating band was caused by protein phosphorylation, we subjected immunoprecipitates of GFP-DSB-1 to λ-phosphatase treatment. Phosphatase treatment abolished the slow-migrating band, showing that this was a phosphorylated fraction of DSB-1 (**Figure 1B**). Taken together, these results strongly suggest that phosphorylated DSB-1 protein is normally dephosphorylated in a PPH-4.1-dependent manner in wild-type animals.

**Figure 1.**
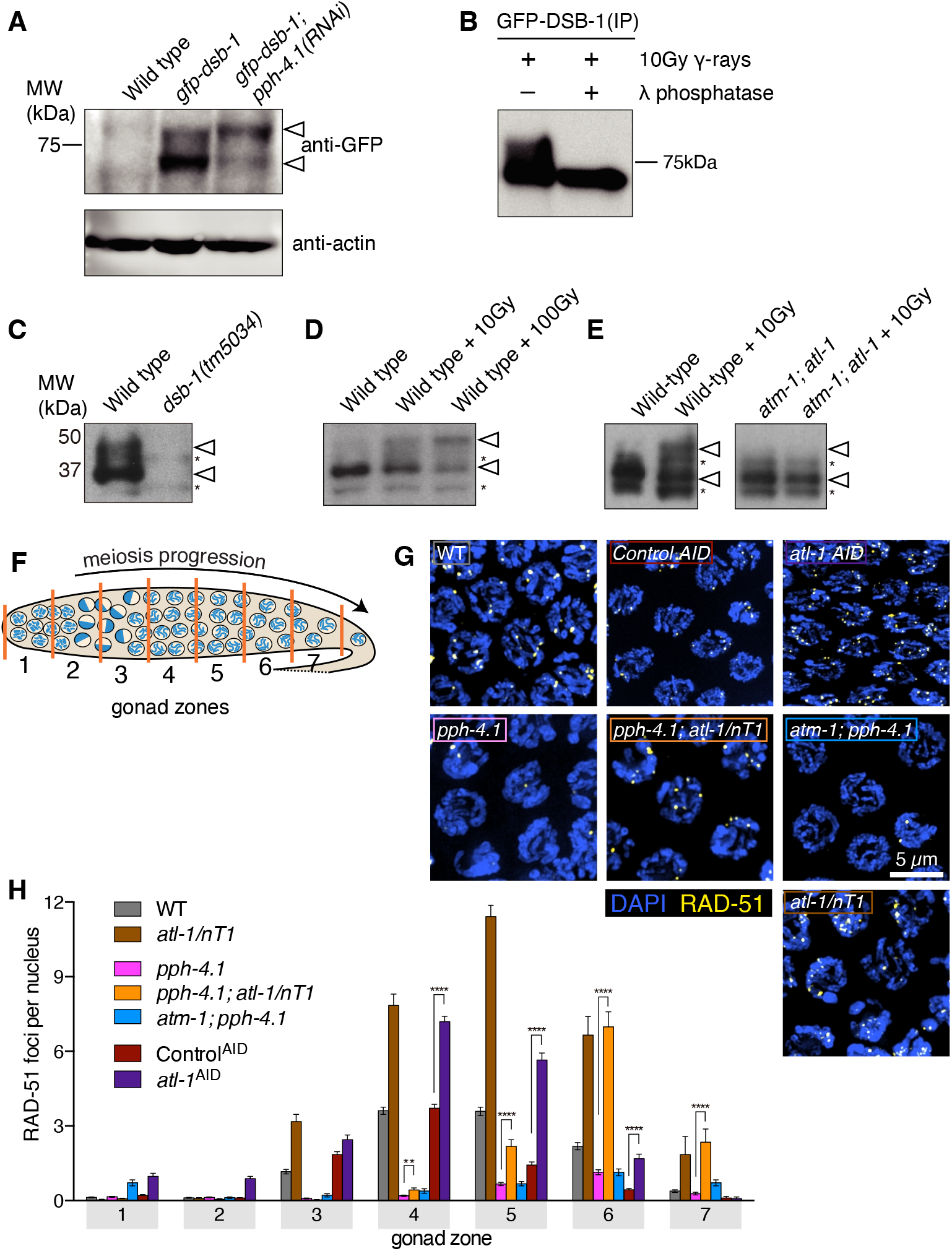
DSB-1 is phosphoregulated in a PPH-4.1^PP4^- and ATL-1^ATR^ -dependent manner and ATL-1^ATR^ kinase antagonizes PPH-4.1^PP4^ phosphatase. **(A)** Western blot of GFP-fused DSB-1 probed with α-GFP. GFP-DSB-1 detected in extracts from wild type and *gfp::dsb-1* worms (24 h post L4 stage) with either control RNAi or *pph-4*.*1* RNAi treatment. A total protein amount of 97 μg was loaded in each lane. Arrowheads indicate two specific bands in the blot. Loading controls (α-actin) are shown at bottom. **( B)** Blot of GFP-DSB-1 purified by immunoprecipitation from young adult RNAi-treated worms subjected to 10Gy γ-irradiation, with or without treatment with λ-phosphatase. **(C, D, E)** Western blots of endogenous DSB-1 probed with α-DSB-1 antibodies in wild type (N2), *dsb-1(tm5034), atm-1(gk186), atl-1(tm853*) or *atm-1(gk186); atl-1(tm853)* with combination of γ-irradiation (10 Gy in panel E, and 10 or 100 Gy in panel D as indicated). Lysate of 50 worms at 24 h post L4 stage was loaded in each lane. Asterisks show non-specific bands, and arrowheads indicate two specific bands. **(F)** Schematic showing a hermaphrodite gonad divided into 7 equally-sized zones for RAD-51 focus scoring. (**G**) Immunofluorescence images of RAD-51 foci in mid-pachytene nuclei (zone 5) of the indicated genotypes. Scale bar, 5 μm. **(H)** Quantification of RAD-51 foci in the germlines of the genotypes indicated in **(G)**. Seven gonads were scored for wild type and *pph-4*.*1(tm1598)*, three gonads were scored for *atl-1(tm853)/nT1, atl-1*^*AID*^, *control*^*AID*^ as well as *pph-4*.*1(tm1598); atl-1(tm853)/nT1* double mutants, and four gonads were scored in *atm-1(gk186); pph-4*.*1(tm1598)*. The numbers of nuclei scored in zones 1-7 were as follows: for wild type, 420, 453, 377, 375, 345, 271, 296; for *pph-4*.*1(tm1598)*, 433, 423, 422, 413, 355, 322, 208; for *atl-1(tm853)/nT1*, 103, 137, 145, 115, 97, 75, 40; for *atl-1*^*AID*^, 161, 193, 180, 241, 204, 117, 37; for *control*^*AID*^, 143, 188, 233, 223, 192, 109, 28; for *pph-4*.*1(tm1598); atl-1(tm853)/nT1*, 126, 121, 98, 100, 94, 86, 49; for *atm-1(gk186);pph-4*.*1(tm1598)*, 123, 153, 167, 140, 161, 156, 123. Significance was assessed via two-tailed *t* test, **P<0.01, ****P<0.0001.

### ATL-1^ATR^ kinase opposes the DSB initiation activity of PPH-4.1^PP4^ phosphatase

To examine if ATM/ATR kinase may phosphorylate DSB-1 and thus antagonize PPH-4.1 phosphatase activity, we performed western blots to detect endogenous DSB-1 in *atm-1*^*ATM*^; *atl-1*^*ATR*^ double mutants in combination with γ-ray irradiation to activate ATM/ATR kinases. A phospho-DSB-1 band visible in wild type animals showed a relatively increased fraction in a manner dependent on γ-ray dose (**Figure 1C, D**). This phosphorylated band was abolished in *atm-1; atl-1* double mutants even after γ-irradiation, suggesting that DSB-1 becomes phosphorylated in an ATM/ATR dependent manner (**Figure 1E**). We also conducted the same γ-ray irradiation in mutants lacking DSB formation (*spo-11(me44), htp-3(tm3655), chk-2(me64), rad-50(ok197), mre-11(ok179)*) (2, 41–44). In all these mutant backgrounds, the phosphorylated band was absent before irradiation. Upon γ-irradiation, the phosphorylated band in *spo-11, htp-3*, and *chk-2* mutants became visible, but was undetectable in *mre-11* mutants and extremely faint in *rad-50* mutants (**Supplemental Figure 1C, D**). This is consistent with the previous observation that the MRN (Mre11/Rad50/Nbs1) complex plays an important role in activating ATM/ATR kinases (45–48). These data further reinforce the hypothesis that DSB-1 phosphorylation depends on DSBs and activation of ATM/ATR kinases.

To further examine if ATM/ATR kinases antagonize PPH-4.1 phosphatase to regulate DSB-1 activity, we combined the *pph-4*.*1(tm1598)* mutation with the *atm-1* or *atl-1* mutations. We assessed DSB formation in these double mutants by performing immunofluorescence against the strand-exchange protein RAD-51. Since the *C. elegans* gonad contains nuclei from all stages of meiotic prophase arranged sequentially along the distal-proximal axis, we divided the gonad into 7 zones of equal size (**Figure 1F**) and counted the number of RAD-51 signals in nuclei within each zone to assess the kinetics of DSB initiation. As shown previously, mutation of *pph-4*.*1* led to a drastic decrease of the number of foci compared to wild-type controls (**Figure 1G, H**). Homozygous null mutation of ATM (*atm-1* in *C. elegans*) in a *pph-4*.*1* background only marginally increased the number of DSBs in very late pachytene. In contrast, we found that heterozygous mutation of ATR (*atl-1* in *C. elegans*) in a *pph-4*.*1* mutant led to a strong recovery of RAD-51 foci (**Figure 1G, H**). These results suggest that ATR kinase normally acts to suppress formation of meiotic DSBs, and this activity is opposed by PPH-4.1. Consistent with this hypothesis, we found that auxin-induced depletion (49) of AID-tagged ATL-1 led to a significant increase in RAD-51 foci compared to auxin-treated controls (**Figure 1H**). Similarly, we found that a heterozygous null *atl-1(tm853)* mutation led to a significant increase in overall DSB number in the presence of wild-type *pph-4*.*1*. (**Figure 1H**). The extra DSBs seen upon loss of ATL-1 were not a result of mitotic DNA damage, since the premeiotic zone in both auxin-depleted and *atl-1* heterozygous germlines is mostly free of RAD-51 foci (**Supplemental Figure 2A**), as is also the case in wild-type. Since homozygous mutation of *atl-1* leads to severe mitotic defects and aneuploidy in the germline due to replication errors (48) (**Supplemental Figure 2B**), the numerous RAD-51 foci in homozygous *atl-1* mutants were not scored. In *C. elegans, atm-1* homozygous null animals derived from heterozygous mothers are superficially wild type and fertile. However, *atm-1* mutants are sensitive to DNA damage-inducing reagents, and when maintained homozygously for more than 20 generations, they develop genomic instability and embryonic inviability (50). To assess ATM-1’s contribution to DSB formation during meiotic prophase, we examined the null allele *atm-1(gk186)* in a *rad-54(ok615)* mutant background, in which DSBs are initiated but RAD-51 cannot be removed from recombination intermediates (51, 52). We found that *atm-1; rad-54* germlines showed a significant delay in RAD-51 loading in early pachytene compared to *rad-54* single mutants but eventually showed a level of foci exceeding that of the control in late pachytene (**Supplemental Figure 2C**). Initial delay in RAD-51 foci appearance is consistent with a previously-described role of ATM-1 in timely loading of RAD-51 (53). Taken together, these results show that ATL-1 plays the major role in antagonizing the DSB-promoting function of PPH-4.1, while the role of ATM-1 is less significant.

### DSB-1 possesses conserved SQ motifs, and its non-phosphorylatable mutants rescue the phenotypes of PP4 mutants

DSB-1 possesses five ATM/ATR consensus motifs ([ST]Q), two of which are highly positionally conserved in the genus *Caenorhabditis* (**Figure 2A, Supplemental Figure 3)**. These SQ sites are dispersed within a large region predicted to be intrinsically disordered (**Figure 2A**). The presence of these consensus motifs raises the possibility that PPH-4.1 may dephosphorylate DSB-1 at one or more of these sites to increase DSB-promoting activity. To examine whether hyperphosphorylation of DSB-1 is responsible for loss of DSB initiation in *pph-4*.*1* mutants, we constructed a nonphosphorylatable mutant allele, *dsb-1(5A)*, in which all five SQ serines were replaced by alanine (**Figure 2A**). This non-phosphorylatable allele was fully viable when homozygous, demonstrating that these substitutions do not compromise the activity of DSB-1 protein (**Table 1**). We then examined RAD-51 foci in *dsb-1(5A); pph-4*.*1* animals to assess DSB formation. Germlines homozygous for both *pph-4*.*1* and *dsb-1(5A)* showed a drastically higher number of RAD-51 foci compared to *pph-4*.*1* single mutants (**Figure 2B, C**). Indeed, single mutants of *dsb-1(5A)* alone showed significantly elevated DSB levels compared to wild-type controls, suggesting that phosphorylation of these serine residues acts in wild-type germlines to limit the number of DSB initiations. We also observe a delayed peak in RAD-51 foci in *pph-4*.*1* and *dsb-1(5A); pph-4*.*1*, as well as in *pph-4*.*1; atl-1/nT1* (**Figure 1H, Figure 2C**). Previous studies have shown that PP4 homologs are involved in multiple steps in the processing of somatic cell recombination intermediates involving resection of DSBs and loading of RAD-51 in yeast and mammals (37, 38, 54). PPH-4.1 may also contribute to timely processing of recombination intermediates in meiosis in a similar manner, leading to a delay in RAD-51 loading in all strains lacking PPH-4.1.

**Figure 2.**
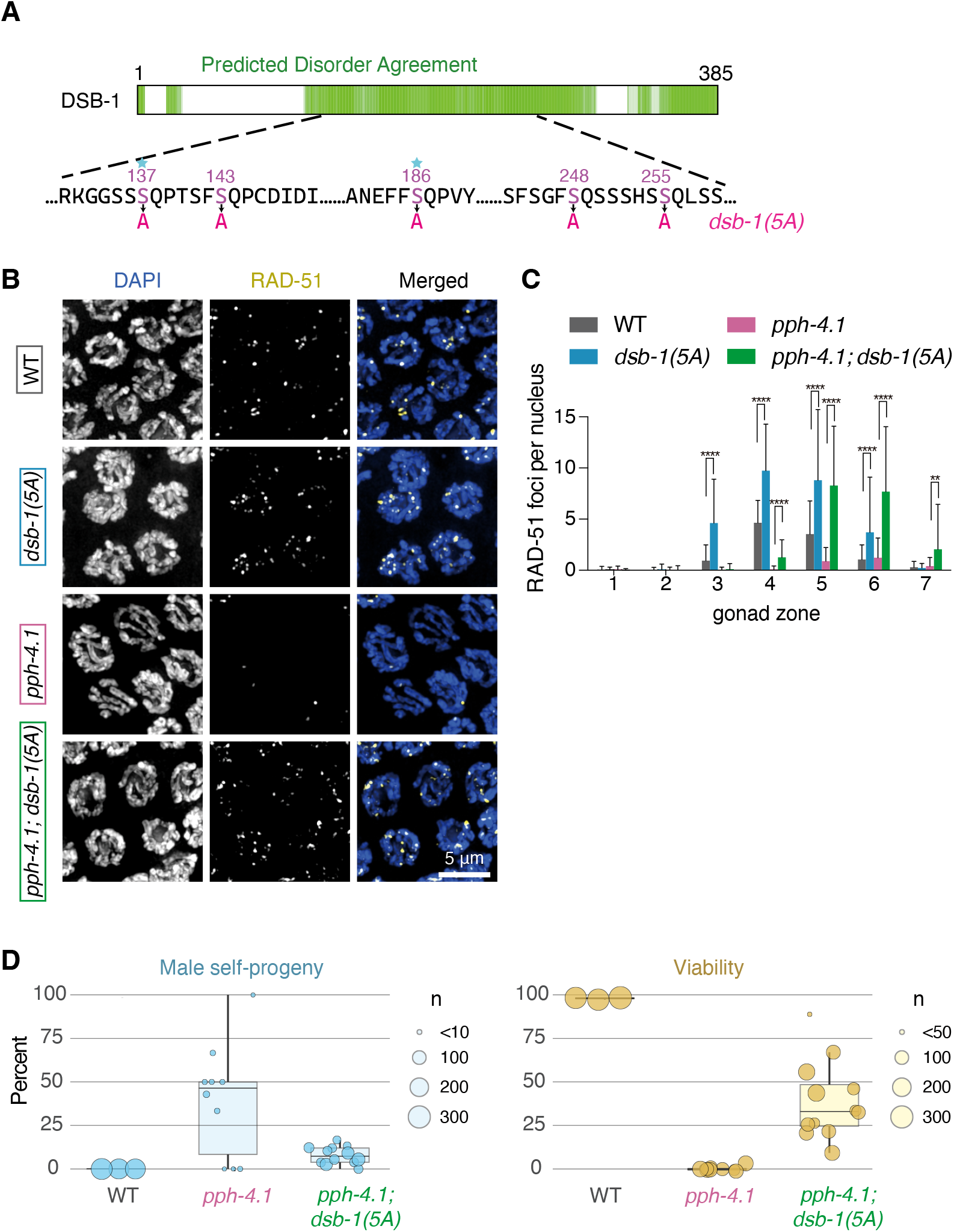
The *dsb-1(5A)* mutation rescues DSB defect and viability loss of *pph-4*.*1* mutants. **(A)** A schematic diagram of the DSB-1 protein sequence. Green regions indicate intrinsically disordered regions from the D^2^P^2^ database (82). Five serines which were mutated into alanines in *dsb-1(5A)* within the SQ sites are shown in magenta, and sites conserved in 10 or more of the 11 Caenorhabditids in the *Elegans* group (see **Supplemental Figure 3**) are indicated with a star. **(B)** Immunofluorescence images of RAD-51 foci in mid-pachytene nuclei of indicated genotypes. Scale bar, 5 μm. **(C)** Quantification of RAD-51 foci in the gonads of indicated genotypes in **(B)**. Four gonads were scored in each genotype; the numbers of nuclei scored in zone 1-7 were as follows: for wild type, 162, 201, 204, 230, 209, 155, 105; for *pph-4*.*1(tm1598)*, 145, 174, 201, 188, 167, 149, 74; for *dsb-1(5A)*, 175, 220, 197, 163, 143, 116, 90; for *pph-4*.*1(tm1598); dsb-1(5A)*, 171, 152, 121, 180, 185, 137, 85. Significance were assessed via the two-tailed *t* tests, **P<0.01, ****P<0.0001. **(D) *Left:*** Male progeny percentage indicating the rate of X chromosome nondisjunction during meiosis in wild type, *pph-4*.*1(tm1598)* and *pph-4*.*1(tm1598); dsb-1(5A)* mutants; circle size corresponds to total number of adult animals scored. ***Right:*** Embryonic viability percentage of the indicated genotypes; circle size corresponds to total number of eggs laid.

**Table 1.** Embryonic viability, male progeny percentage indicating the rate of X chromosome nondisjunction, and total number of scored embryos is shown for hermaphrodite self-progeny of the indicated genotypes.

| Genotype | Embryonic viability (%) | Male percentage (%) | Total # eggs scored |
| --- | --- | --- | --- |
| WT | 99.28 | 0.04 | 1990 |
| *gfp-dsb-1* | 99.10 | 0.19 | 2578 |
| *dsb-1(1A)* | 98.47 | 0.15 | 3427 |
| *dsb-1(2A)* | 98.25 | 0.00 | 1347 |
| *dsb-1(3A)* | 99.29 | 0.19 | 2056 |
| *dsb-1(5A)* | 99.22 | 0.00 | 2546 |
| *pph-4.1(tm1598)* | 2.00 | 45.60 | 1424 |
| *dsb-2(me96)* | 39.55 | 13.85 | 3413 |
| *dsb-2(me96);dsb-1(1A)* | 93.96 | 1.40 | 3370 |
| *dsb-2(me96);dsb-1(2A)* | 83.12 | 3.31 | 2413 |
| *dsb-2(me96);dsb-1(3A)* | 97.56 | 0.64 | 2395 |
| *dsb-2(me96);dsb-1(5A)* | 98.64 | 0.04 | 1596 |
| *pph-4.1(tm1598);dsb-1(1A)* | 7.39 | 32.21 | 496 |
| *pph-4.1(tm1598);dsb-1(5A)* | 40.89 | 7.89 | 1273 |

We next quantitatively scored the meiotic competence and embryonic viability of *dsb-1(5A); pph-4*.*1* double mutants. Since meiotic nondisjunction of the X chromosome leads to production of males (with a single X chromosome) among the self-progeny of *C. elegans* hermaphrodites, we examined the frequency of male progeny in *pph-4*.*1; dsb-1(5A)* mutants, and found that it decreased significantly compared to *pph-4*.*1* single mutants (**Figure 2D, left**). While *pph-4*.*1* single mutants have a very low embryonic viability of 2% on average, the viability of *pph-4*.*1*; *dsb-1(5A)* double mutants increased to 41% (**Figure 2D, right; Table 1**). Taken together, our results strongly suggest that DSB-1 is dephosphorylated in a PPH-4.1-dependent manner to promote DSB formation.

### DSB-1 non-phosphorylatable mutants rescue the homologous pairing defect of PP4 mutants

The striking increase in viability we observed in *pph-4*.*1; dsb-1(5A)* animals was somewhat surprising, given our previous finding that autosomal pairing and synapsis is severely reduced in *pph-4*.*1* single mutants (40). Pairing of an autosome (chromosome V) did not exceed 25% in *pph-4*.*1* mutants, making the expected probability for all five autosomes to pair homologously less than 0.1%, assuming similar rates of pairing for the other autosomes. Actual measurements of synapsis and bivalent numbers at diakinesis in *pph-4*.*1* mutants support this expectation, since formation of six bivalents as in wild type worms is extremely rarely seen (40). We therefore hypothesized that the elevated DSB levels in *pph-4*.*1; dsb-1(5A)* mutants might promote bivalent formation by increasing the level of homologous pairing. To test this, we carried out fluorescence *in situ* hybridization (FISH) against the 5S rDNA locus on chromosome V to assess the progression of homologous pairing over time, in *pph-4*.*1* compared to *pph-4*.*1; dsb-1(5A)* mutants. While the *pph-4*.*1* mutant showed very low homologous pairing as expected, the *pph-4*.*1; dsb-1(5A)* double mutant rescued pairing to a significant degree, up to 70% (**Figure 3A, B**). Immunostaining against the protein ZIM-3, which marks the pairing center end of chromosomes I and IV (55), in *pph-4*.*1; dsb-1(5A)* and *pph-4*.*1*, gave qualitatively similar results (**Supplemental Figure 4A**). Consistent with recent discoveries that synapsis prior to recombination in *C. elegans* is a dynamic state that later becomes stabilized by recombination (33–35, 56, 57), these results indicate that introduction of DSBs into a *pph-4*.*1* mutant also leads to increased fidelity of pairing and synapsis.

**Figure 3.**
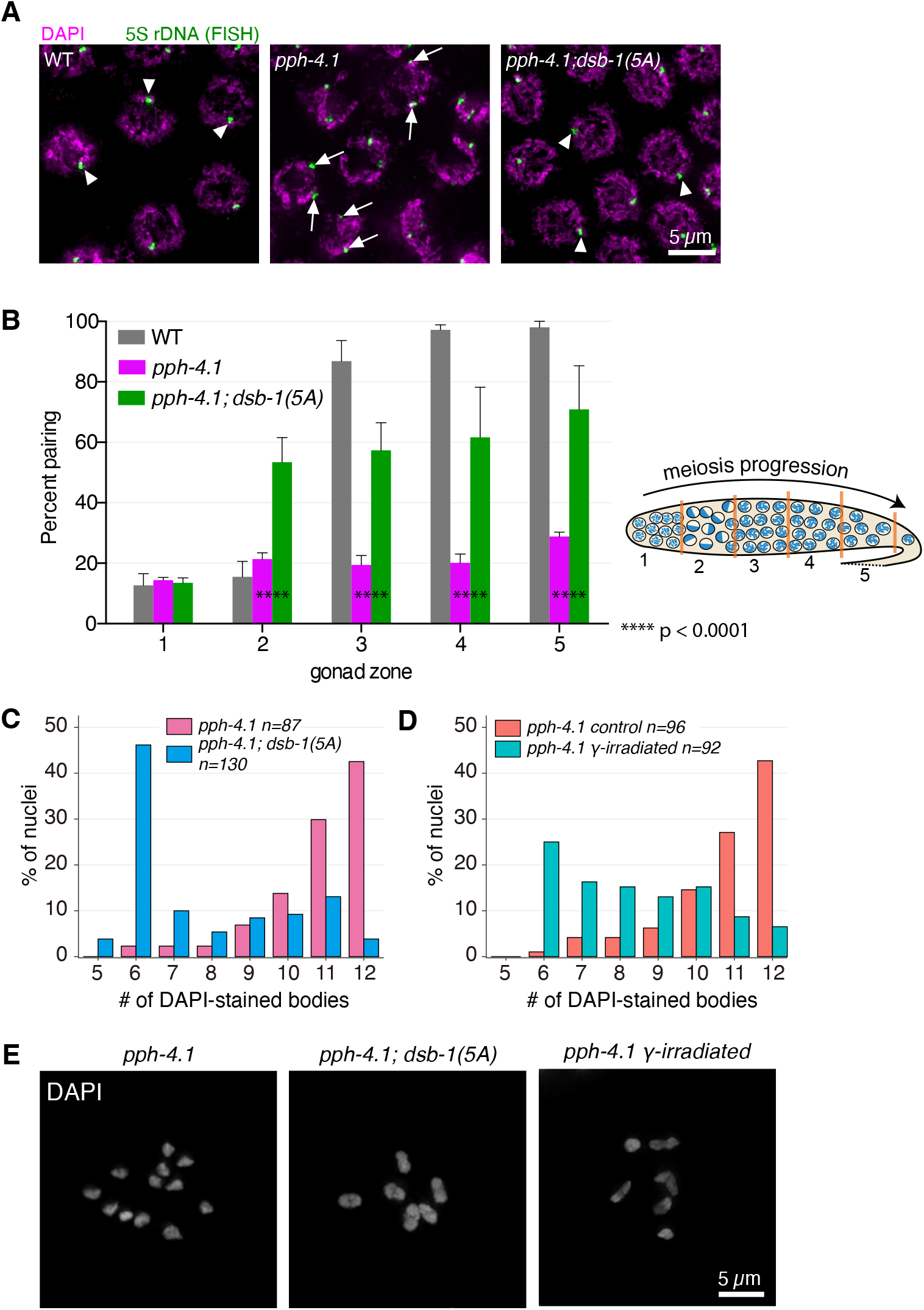
Chiasma formation and homologous pairing defects are partially rescued by the *dsb-1(5A)* allele in *pph-4*.*1* mutants. **(A)** FISH images show paired 5S rDNA sites in wild type and *pph-4*.*1(tm1598); dsb-1(5A)* worms (left and right; arrowheads indicate paired foci), and unpaired sites in *pph-4*.*1(tm1598)* mutants (middle; arrows indicate unpaired foci) at pachytene. Scale bar, 5 μm. **(B) *Left:*** Quantification of pairing for chromosome V shown as the percent of nuclei with paired signals in each zone. Four gonads were scored for each genotype. The total number of nuclei scored for zone 1-5 respectively was as follows: for wild type, 163, 266, 208, 148, 76; for *pph-4*.*1(tm1598)*, 300, 283, 257, 219, 124; for *pph-4*.*1(tm1598); dsb-1(5A)*, 335, 318, 266, 224, 137. Significance was assessed by chi-squared test for independence, ****P<0.0001. ***Right:*** Schematic showing a hermaphrodite gonad divided into 5 equally-sized zones for FISH focus scoring. **(C)** The number of DAPI-stained bodies shown as percentages of the indicated number of diakinesis oocyte nuclei scored for *pph-4*.*1(tm1598)* and *pph-4*.*1(tm1598); dsb-1(5A)* mutants. The numbers of nuclei scored for each genotype were: 87 for *pph-4*.*1(tm1598)*, 130 for *pph-4*.*1(tm1598); dsb-1(5A)*. **(D)** The number of DAPI-stained bodies shown as percentages of the indicated number of diakinesis oocyte nuclei scored in *pph-4*.*1(tm1598)* mutants with or without γ-irradiation. The numbers of nuclei scored for each genotype were: 96 for *pph-4*.*1(tm1598) control*, 92 for *pph-4*.*1(tm1598)* γ*-irradiated*. **(E)** Images of DAPI-stained diakinesis nuclei in a *pph-4*.*1(tm1598)* mutant and a *pph-4*.*1(tm1598); dsb-1(5A)* double mutant, as well as a *pph-4*.*1(tm1598)* mutant exposed to 50 Gy of γ-irradiation. Scale bar, 5 μm.

We verified the formation of bivalents in *pph-4*.*1;dsb-1(5A)* animals by counting the number of DAPI-stained bodies in oocytes at diakinesis (**Figure 3C, E**). In the wild type, nearly 100% of diakinesis nuclei show six DAPI-stained bodies corresponding to six bivalents whereas most of the nuclei carry univalents in *pph-4*.*1* single mutants (40). As expected from the increase in homolog pairing, the number of bivalents in *pph-4*.*1; dsb-1(5A)* mutants was significantly higher than in *pph-4*.*1* single mutants.

The apparent rescue of embryonic viability by increased DSBs in *pph-4*.*1; dsb-1(5A)* mutants seemed contradictory to our previous observation that γ-ray irradiation (10 Gy) did not rescue bivalent formation in *pph-4*.*1* mutants (40). However, at a higher dose of γ-irradiation (50 Gy), we were able to rescue bivalent formation in *pph-4*.*1* mutants, with 25% of oocytes showing 6 bivalents (**Figure 3D, E**). Since 10 Gy irradiation at least partially rescues bivalent formation in mutants carrying the null *spo-11(me44)* allele but not in *pph-4*.*1* mutants (2, 40), our observation suggests that conversion from DSBs to crossovers is not as efficient in *pph-4*.*1* mutants, and thus *pph-4*.*1* mutants require more DSBs than *spo-11* mutants for bivalent formation. This requirement for higher break numbers is likely imposed by the combined deficiencies of *pph-4*.*1* mutants both in timely processing of recombination intermediates as well as in preventing non-homologous synapsis.

### DSB-1 non-phosphorylatable mutants rescue the defects of *dsb-2* mutants

Many species in the *Caenorhabditis* genus possess a paralog of *dsb-1*, called *dsb-2*. In *C. elegans dsb-2(me96)* null mutants, a profound reduction of DSBs leads to crossover defects, causing severe embryonic inviability that increases with maternal age (24). However, DSBs are not completely eliminated in *dsb-2* mutants, suggesting that while DSB-1 activity in the absence of DSB-2 can suffice to initiate DSBs, this activity is lower compared to when DSB-2 is present. The ability of the non-phosphorylatable *dsb-1(5A)* allele to rescue the loss of DSBs in *pph-4*.*1* mutants suggests that the 5A allele is hyperactive and not subject to downregulation through phosphorylation. To test whether this hyperactive allele depends on DSB-2 for its high levels of break formation, we performed RAD-51 immunofluorescence on *dsb-1(5A); dsb-2* germlines. In agreement with previous observations, we found very few RAD-51 foci in *dsb-2* mutant nuclei. However, the *dsb-1(5A)* allele strongly rescued DSB formation in the *dsb-2* null background to levels higher than wild-type, similar to what is seen in the *dsb-1(5A)* allele alone (**Figure 4A, B**). Thus, DSB formation in the presence of the 5A allele of *dsb-1* appears to be completely insensitive to loss of *dsb-2*, providing further evidence of the allele’s hyperactivity. Moreover, this increased number of DSBs leads to normal bivalent formation and production of fully viable embryos in *dsb-2; dsb-1(5A)* double mutants (**Figure 4C, Table 1**).

**Figure 4.**
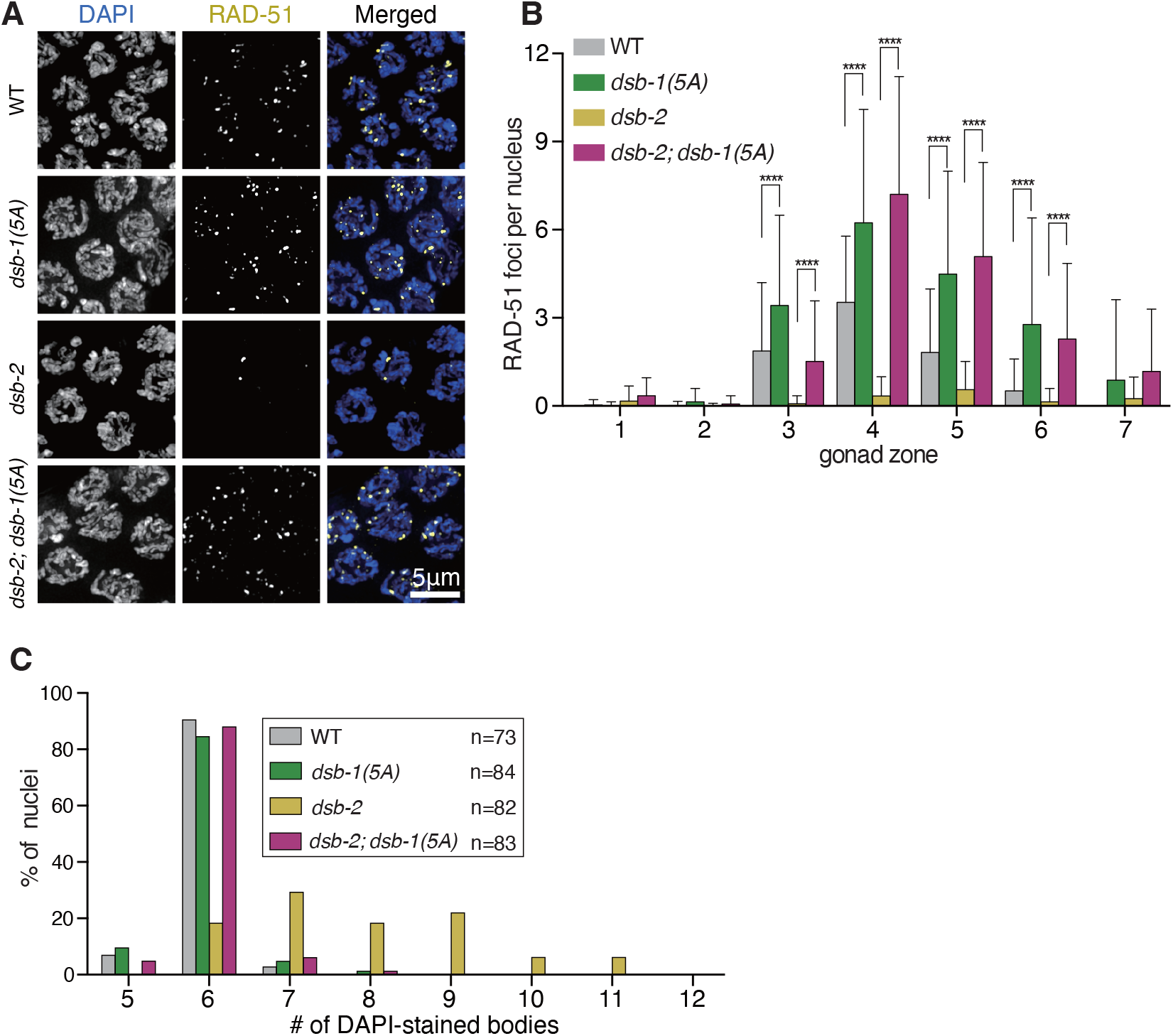
The *dsb-1(5A)* mutation rescues DSB and crossover formation in *dsb-2* mutants. **(A)** Immunofluorescence images of RAD-51 foci in mid-pachytene nuclei of the indicated genotypes. Scale bar, 5 μm. **(B)** Quantification of RAD-51 foci in the gonads of the genotypes indicated in **(A)**. Three gonads were scored in *dsb-2(me96), dsb1(5A)* and *dsb-2(me96); dsb-1(5A)* respectively, and two gonads were scored in wild type; the numbers of nuclei scored in zone 1-7 were as follows: for wild type, 57, 99, 112, 123, 109, 62, 27; for *dsb-2(me96)*, 124, 105, 108, 114, 108, 84, 53; for *dsb-1(5A)*, 126, 119, 103, 118, 116, 101, 79; for *dsb-2(me96); dsb-1(5A)*, 94, 131, 118, 91, 93, 96, 71. Significance was assessed via two-tailed *t* test, ****P<0.0001. **(C)** The number of DAPI-stained bodies shown as percentages of the indicated number of diakinesis oocyte nuclei scored for each genotype. The numbers of nuclei scored for each genotype were: 73 for wild type, 84 for *dsb-1(5A)*, 82 for *dsb-2(me96)*, 83 for *dsb-2(me96); dsb-1(5A)*.

To gain further insight into the five SQ sites, we generated a series of *dsb-1* non-phosphorylatable mutants: *dsb-1(1A)*, which is *dsb-1(S186A)*; *dsb-1(2A)*, which is *dsb-1(S137A_S143A)* and *dsb-1(3A)*, which is *dsb-1(S137A_S143A_S186A)* based on the observation that S137 and S186 are highly conserved within the *Elegans* group of Caenorhabditids (**Figure 5A, Supplemental Figure 3**), and examined if they rescue *dsb-2* mutants. We found that the both *dsb-1(1A)* and *dsb-1(3A)* alleles were sufficient to rescue both embryonic viability and male frequency to wild-type levels at all maternal ages (**Figure 5B, Table 1**). In contrast, the *dsb-1(2A)* mutation, which does not include S186, rescued the embryonic viability and male frequency defects in young *dsb-2* adults (day 1 post-L4 larval stage), but the rescue was less pronounced in older animals (day 3 post -L4) (**Figure 5B**). We therefore conclude that phosphorylation of any of the SQ motifs in the intrinsically-disordered region of DSB-1 may contribute to shutting down the DSB-promoting activity of DSB-1, with S186 phosphorylation likely to be a strong determinant of reduced DSB activity in aged animals.

**Figure 5.**
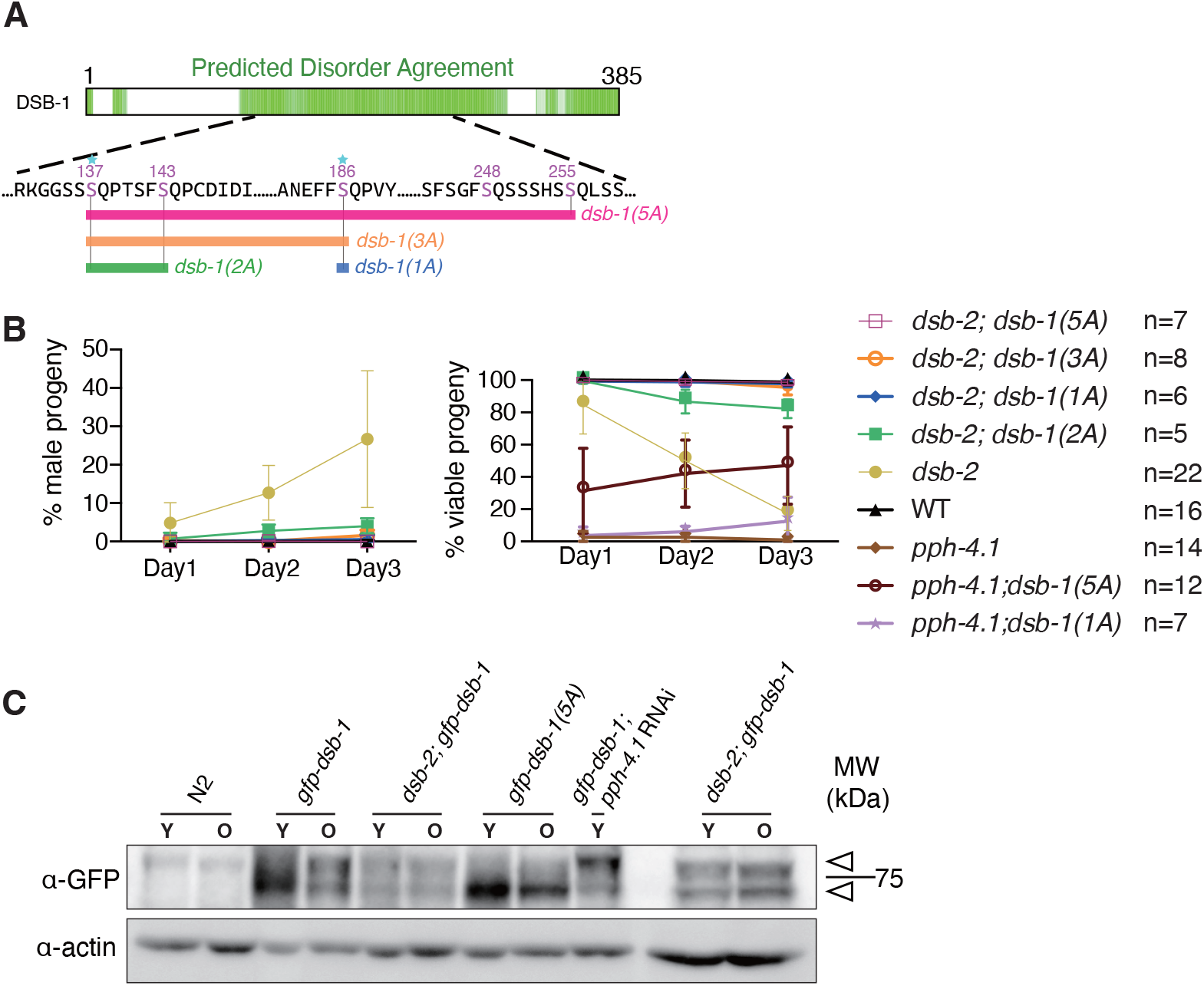
Alanine substitution of serine 186 in DSB-1 suffices to rescue the *dsb-2* mutation. **(A)** Diagram depicting a series of *dsb-1* phospho mutants: *dsb-1(1A)* is *dsb-1(S186A)*; *dsb-1(2A)* is *dsb-1(S137A; S143A)*; *dsb-1(3A)* is *dsb-1(S137A; S143A; S186A)* and *dsb-1(5A)* is *dsb-1(S137A; S143A; S186A; S248A; S255A)*. **(B)** The frequency of males among surviving progeny (left) and the frequency of viable embryos (right) from eggs laid by hermaphrodites of the indicated genotypes during the indicated time interval after the L4 larval stage. **(C)** Western blots of GFP-fused DSB-1 from young adults (Y, 24 h post-L4 larval stage) and old adults (O, 72h post-L4 larval stage) of the indicated genotypes, probed with α-GFP; arrowheads indicate the two GFP-DSB-1-specific bands in the blot. A total protein amount of 97 μg was loaded in each lane except for the two lanes of *dsb-2; gfp-dsb-1* double mutants on the right, in which 162 μg protein was loaded in each lane. Loading controls (α-actin) are shown at bottom.

While the *dsb-1(1A)* allele fully rescued *dsb-2* mutants, it did not rescue embryonic viability in *pph-4*.*1* mutants (**Figure 5B, right; Table 1**). Gonads of *dsb-1(1A)* mutants showed comparable levels of RAD-51 foci to wild type, and introducing the *dsb-1(1A)* mutation increased RAD-51 foci both in *pph-4*.*1; dsb-1(1A)* and *dsb-2; dsb-1(1A)* mutants compared to respective single mutant, *pph-4*.*1* or *dsb-2* (**Supplemental Figure 4B**). The difference in embryonic viability likely reflects inefficient processing of DSBs to COs in *pph-4* mutants, similar to the prior observation that higher levels of gamma irradiation was required to rescue bivalent formation in this mutant.

We and others have shown that DSB initiation activity is significantly reduced in older adults (at least 72 h post-L4) compared to young adults (24 h post-L4) using *rad-54(ok617)* mutants, in which DSBs are generated but the resulting recombination intermediates are trapped without completion of repair (40, 58). We therefore wondered whether DSB-1 may become more phosphorylated with age, thereby contributing to age-dependent reduction of DSB activity. To test this, we performed western blotting to assess DSB-1 phosphorylation levels in young (24 h post-L4) versus older (72 h post-L4) animals. Protein extracts were made from either wild-type, *gfp*-*dsb-1* or *gfp*-*dsb-1(5A)* worms in varying combination with *dsb-2* and *pph-4*.*1* RNAi, blotted onto membranes and probed with α-GFP antibodies. In *gfp*-*dsb-1* animals, the proportion of a slow-migrating band increased in older animals (**Figure 5C**), suggesting an increased proportion of phosphorylated DSB-1. An overall reduced amount of GFP-DSB-1 was detected in the *dsb-2* background, consistent with a previous study (23). The phosphorylated band was proportionally stronger in young *dsb-2* mutants compared to young control animals, suggesting that both reduced protein levels and increased phosphorylation on DSB-1 likely contribute to lower DSB activity in *dsb-2* young animals (**Figure 5C**). However, somewhat surprisingly, the proportion of phosphorylated versus unphosphorylated GFP-DSB-1 did not change with age in *dsb-2* mutants. We verified this by loading a higher amount of protein and comparing the density of phosphorylated versus unphosphorylated bands in *gfp-dsb-1; dsb-2* (rightmost two lanes of **Figure 5C**). This raises the possibility that something other than DSB-1 phosphorylation may contribute to age-dependent degradation of DSB production in *dsb-2* older animals. Alternatively, phosphorylation specifically on S186 may increase with age in *dsb-2* mutants, but this change may not be not detectable without S186 phos-specific antibodies. We attempted to generate specific antibodies against DSB-1 phospho-S186 twice, but failed to obtain a specifically-staining antibody. In *gfp-dsb-1(5A)* animals, the fraction of phosphorylated bands is reduced both in young and old animals compared to the control. While the slow-migrating band is absent, a smearing of GFP-DSB-1(5A) can be seen above the main band, suggesting that some other of the 86 serines, threonines, or tyrosines in this protein may still be phosphorylated in the absence of phosphorylation at the five SQ sites (**Figure 5C**). Taken together, our results show age-dependent increase of DSB-1 phosphorylation likely contributes to a reduction in DSB levels in wild type.

## Discussion

In this work, we have identified DSB-1 as a nexus of DSB initiation control by phosphoregulation in *C. elegans*. While negative regulation of the Spo11 cofactor Rec114 by the DNA damage kinases ATM and ATR has been previously investigated (11), we show here for the first time that PP4 phosphatase plays an opposing role, working through DSB-1 to promote a number of breaks sufficient to generate a crossover on every chromosome. Moreover, we show that ATR kinase plays a key inhibitory role in *C. elegans* DSB initiation. Based on our results, the most straightforward model is that PPH-4.1 dephosphorylates DSB-1 to promote DSBs, whereas ATL-1 phosphorylates DSB-1 to prevent excess DSB production, and this phospho-regulation circuit ensures termination of DSB production only after sufficient numbers of DSBs are generated. This is consistent with the previous observations that many ATM/ATR targets such as Hop1, Mek1, RPA2, H2A and Zip1 are dephosphorylated in a PP4 dependent manner in other organisms (36, 37, 59, 60). However, since other factors including an effector kinase Rad53(Chk2) are also known to be dephosphorylated in a PP4-dependent manner in budding yeast (Villoria et al. 2019), we note that we cannot exclude the possibility that PPH-4.1 indirectly reduces DSB-1 phosphorylation by dephosphorylating upstream factors regulating DSB-1.

Since the mechanism by which the RMM group of cofactors (including DSB-1/2) promotes Spo11 activity remains unknown, how phosphorylation of these factors negates their activity is also not clear. Recent work has shown that neutralizing a conserved basic patch of either Rec114 or Mer2 leads to loss of DNA binding (26); analogously, DSB-1 interaction with DNA may be prevented by the negative charge of phosphorylation. The same work showed that RMM proteins form DNA-dependent condensates, which raises the possibility that phosphorylation of the intrinsically disordered region could electrostatically alter any phase separation propensity of DSB-1. Further investigation of DSB-1 and DSB-2 *in vivo* is necessary to determine the extent to which they resemble their yeast orthologs with regard to condensate formation.

The recent identification of *dsb-3* as a nematode ortholog of Mei4, and its likely participation in a complex with DSB-1 and DSB-2 (25) akin to the heterotrimeric Rec114-Mei4 complex of yeast (26) prompted us to examine predicted structural properties of such a complex using the AlphaFold structure prediction pipeline (61, 62). Since both DSB-1 and DSB-2 are orthologs of Rec114, we tested three possible complexes: a fully heterotrimeric complex of DSB-1, DSB-2, and DSB-3, and complexes containing two copies of either DSB-1 or DSB-2 with one copy of DSB-3. We also generated predictions for the DSB-1,-2, and -3 orthologs from the closely related species *Caenorhabditis inopinata*, as well as for human and yeast Rec114-Mei4 complexes in 2:1 stoichiometry. In the predicted DSB-1:2:3 trimer, the alpha-helical C-termini of DSB-1 and DSB-2 wrap around each other to form a channel which accommodates the helical N-terminus of DSB-3; a similar structure was predicted for a trimer of two Rec114 and one Mei4 (**Supplemental Figure 5A**). In all species (nematode, yeast, and human), these structural predictions are in agreement with previous models based on yeast two-hybrid (25, 63) and crosslinking mass spectrometry analysis (26). This trimer prediction was not found in models of a DSB-2:DSB-2:DSB-3 trimer, but was found in three out of five DSB-1:DSB-1:DSB-3 models. These models raise the possibility that DSB-1 dimers are more likely to bind DSB-3 in the absence of DSB-2, than DSB-2 dimers are to bind DSB-3 in the absence of DSB-1. This asymmetry would be consistent with the more severe phenotype of *dsb-1* compared to *dsb-2* mutants, as well as with yeast two-hybrid evidence showing DSB-1 but not DSB-2 directly binds to DSB-3 (25). The consistency of the structural prediction in three highly-diverged species, and its agreement with known *in vivo* data, is highly suggestive of a conserved interaction. However, since the interacting regions within the predicted trimer does not involve the disordered domain of DSB-1 or any of its SQ motifs, this prediction does not suggest that phosphorylation of DSB-1 would interfere with its ability to bind to the other members of this complex. DSB-2 also possesses four potential ATM/ATR phosphorylation sites (SQs) in its predicted disordered region, and it remains to be seen whether these sites are subject to phosphorylation or if DSB-2 is refractory to phosphoregulation.

The diverged functions of DSB-1 and DSB-2 raise the question of which one of the paralogs is closer to the ancestral gene. We constructed a phylogenetic tree based on the longest isoform of each set of genes and found that the duplication of a single Rec114 ortholog into the paralogs DSB-1 and DSB-2 occurred early within the genus Caenorhabditis. Based on our inferred gene tree, we estimate that the *C. elegans* DSB-1 and DSB-2 protein sequences have undergone 1.36 and 2.02 amino acid substitutions per site, respectively, since the gene duplication event (**Supplemental Figure 5B**). The mean number for all DSB-1 and DSB-2 orthologs since the gene duplication event is 1.03 and 1.66 amino acid substitutions per site, respectively. This suggests that DSB-1 retains the essential ancestral function common to all Rec114 orthologs, while DSB-2 has evolved to perform a slightly modified role. We have demonstrated an increase in the proportion of phosphorylated compared to unphosphorylated DSB-1 in *dsb-2* young animals (**Figure 5C**), and showed that the hyperactive *dsb-1(S186A)* mutant rescues loss of *dsb-2*. These results suggest that DSB-2 is required to counteract the phosphorylation and thus downregulation of DSB-1. Taken together with previous results showing that loss of *dsb-2* leads to reduction in DSB-1 protein levels (23, 24), it is likely that DSB-2 has evolved to perform an auxiliary role in DSB formation, compensating for DSB-1’s gradual deactivation, and thereby extending the window of fertility. We hypothesize that either increasing phosphorylation of DSB-1 S186 with age, or age-correlated increase of another factor that impedes DSB formation by phosphorylated DSB-1, underlies this drop in DSB formation with age seen in *C. elegans*.

Our observation that *dsb-1(5A)* increases the amount of homologous pairing in *pph-4*.*1* mutants (**Figure 3A, B; Supplemental Figure 4A**) adds to growing evidence that while initial pairing and synapsis in *C. elegans* does not depend on DSBs, stabilization and (when necessary) correction of synapsis does. The promiscuous pairing and synapsis in *pph-4*.*1* mutants cannot be attributed solely to a low number of DSBs, since null mutants of both *spo-11* and *dsb-1* completely lacking in DSBs, as well as *dsb-2* mutants with severely reduced DSBs, synapse homologously (2, 23, 24). We hypothesize that in the absence of PP4 activity, hyperphosphorylation of one or more additional substrates leads to promiscuous synapsis in a low-DSB (i.e., immature SC) environment, but providing additional DSBs enforces a higher degree of synaptic fidelity, perhaps via homologous recombination. Further, the number of DSBs required to rescue embryonic viability in *pph-4*.*1* mutants must be higher than that needed to rescue viability in *dsb-2*, since the *dsb-1(1A)* allele suffices to fully rescue mutations in *dsb-2*, but has little effect on *pph-4*.*1* (**Figure 5B, Table 1**). This inconsistency may be resolved by noting that rescue of viability in *pph-4*.*1* mutants requires rescue of both promiscuous pairing and of crossover failure; since *pph-4*.*1; dsb-1(1A)* germlines show numbers of RAD-51 foci intermediate between *pph-4*.*1* and *pph-4*.*1; dsb-1(5A)* (**Supplemental Figure 4B, Figure 2B**), we hypothesize that the number of DSBs required to correct promiscuous pairing in *pph-4*.*1* mutants is higher than that needed to guarantee a crossover on each chromosome. Incomplete rescue of homologous pairing, and incompletely penetrant phenotypes in other known PP4-dependent processes such as centrosome maturation and sperm production (64, 65) that are not rescued by DSBs are likely responsible for the remaining brood size and viability defects in *pph-4*.*1; dsb-1(5A)* double mutants (**Figure 2D**). The fact that worms carrying the *dsb-1(5A)* mutation enjoy full viability, with brood size and male incidence nearly identical to wild-type despite the roughly twofold higher number of RAD-51 foci observed, raises the question of what role negative control of DSBs is playing in *C. elegans*. To examine whether *dsb-1(5A)* could sensitize worms to external DNA damage, we have γ-irradiated control and *dsb-1(5A)* animals and assayed for embryonic inviability as an indicator of unrepaired DNA breaks. However, *dsb-1(5A)* mutants showed no difference from control animals in embryonic viability or brood size after 30 or 75 Gy irradiation (data not shown), suggesting that meiocytes have a large capacity to repair DSBs in excess over wild type levels. A recent study examining the effects of loss of germline ATM in mice (66) discovered an increased incidence of large deletions and other rearrangements at hotspots. Limiting the number of DSBs through DSB-1 phosphorylation could forestall such mutagenic events and maintain genome integrity over generations. Further experiments analyzing long-term genome integrity in non-phosphorylatable *dsb-1* mutants are needed to address this issue. While the physiological role of DSB-1 phosphorylation remains unclear, our work provides the first evidence that the ability of PPH-4.1 to regulate DSB-1 phosphorylation levels in meiotic prophase is critical to provide a sufficient number of breaks to ensure chiasma formation on each chromosome pair.

## Materials and Methods

### Worm strains and antibodies

*C*.*elegans* strains were maintained at 20 °C on nematode growth medium (NGM) plates seeded with OP50 bacteria under standard conditions (67). Bristol N2 was used as the wild type strain and all mutants were derived from an N2 background. A list of all strains and antibodies used is provided in an supplemental spreadsheet. All cytological experiments were performed on adult hermaphrodite germlines.

### Generation of mutants via CRISPR-Cas9 genome editing system

CRISPR-Cas9 genome editing using *dpy-10* as co-CRISPR marker (68) was applied to generate *dsb-1* N-terminal tagged (*gfp-dsb-1*) and *dsb-1* non-phosphorylatable lines. A 10 μL mixture containing 17.5 μM trans-activating CRISPR RNA (tracrRNA)/crRNA oligonucleotides (targeting *dsb-1* and *dpy-10*) purchased from Integrated DNA Technologies (IDT, Coralville, Iowa, USA), 17.5 μM Cas9 protein produced by the MacroLab at UC Berkeley, and 6 μM singlestranded DNA oligonucleotide purchased from IDT or 150 ng/μL double-stranded DNA generated from PCR as a repair template was injected into the gonads of 24 h post-L4 larval stage N2 hermaphrodites. To prevent re-editing by the CRISPR-Cas9 machinery, silent mutations were introduced into the target gene *dsb-1*. For *gfp-dsb-1*, an additional linker sequence of 3x glycine was introduced between the target site and GFP-tag sequence. Dpy or Rol F1 animals (*dpy-10* mutation homozygous or heterozygous, respectively) were picked to individual plates to self-propagate overnight and then screened for successful edits by PCR and DNA sequencing. A list of oligonucleotides used is provided in SI Appendix.

### RNA interference

RNAi was carried out by feeding N2 or *gfp-dsb-1* worms with the HT115 bacteria expressing either the empty RNAi vector L4440 obtained from the Ahringer Lab RNAi library (69) or a *pph-4*.*1* RNAi plasmid (40). Worms were first synchronized through starvation and grown to the L4 larval stage on new NGM plates with OP50 bacteria. L4 worms were collected in M9 (41 mM Na2HPO4, 15 mM KH2PO4, 8.6 mM NaCl, 19 mM NH4Cl) + 0.01% Tween buffer, washed three times with M9 buffer and distributed to RNAi plates. About 30 h later, the worms became gravid and were harvested in M9 + 0.01% Tween buffer, washed three times with M9 buffer and bleached for no more than 3 min to obtain F1 embryos. Collected embryos were placed to fresh RNAi plates and grown until 24 h or 72 h after the L4 larval stage. For the immunoprecipitation experiment, worms were harvested in M9 + 0.01% Tween buffer and exposed to 10 Gy γ-irradiation, and then washed two times with M9 buffer. Lastly, the worms were washed with RIPA buffer (150 mM NaCl, 1% Triton X-100, 0.5% Sodium deoxycholate, 0.1% SDS, 50 mM TrisHCl pH 8.0) + 1mM PMSF +1mM DTT and the supernatant was poured off until 2x pellet volume. Worm pellet suspensions were then supplemented with 2x protease inhibitor cocktail + 3mM NaF, frozen in liquid nitrogen and stored at -80 °C. For western blot analysis, worms were harvested and washed three times in M9 buffer and frozen in liquid nitrogen.

### Auxin-induced protein depletion in worms

Depletion of AID-tagged proteins in the *C. elegans* germline was performed as previously described (49). Briefly, 1mM auxin (IAA, Alfa Aesar #10556, Haverhill, Massachusetts, USA) was added into the NGM agar just before pouring plates. *E. coli* OP50 bacteria cultured overnight were concentrated, supplemented with 1mM auxin and spread on plates. These auxin plates were stored at 4 °C in the dark and used within a month. NGM plates supplemented with the solvent ethanol (0.25% v/v) were used as control. To obtain synchronized worms, L4 hermaphrodites were picked from the maintenance plates. Auxin treatment was performed by transferring worms to auxin plates and incubating for the indicated time at 20 °C. Young adult animals (24 h post-L4) were dissected for immunofluorescence analyses.

### Immunoprecipitation and phosphatase assay

Immunoprecipitation of GFP-tagged DSB-1 was performed with the same amount of lysate determined by BCA kit (Pierce BCA protein assay kit #23225, Thermo Scientific, Waltham, Massachusetts, USA) from wild type and *pph-4*.*1* RNAi animals. Protein samples were incubated with GFP-Trap (ChromoTek #gtma-20, Planegg, Germany) at 4 °C overnight with gentle rotation and the collected beads were washed three times with wash buffer (10 mM Tris/Cl pH 7.5, 150 mM NaCl, 0.05 % Nonidet P-40, 0.5 mM EDTA). The beads to which GFP-DSB-1 had been immobilized were subject to the phosphatase assay (NEB lambda phosphatase #P0753S, Ipswich, Massachusetts, USA) following manufacturer’s instruction at 30 °C for 2 h. Then the protein was eluted in the SDS-PAGE sample buffer by boiling at 95 °C for 10 min and used for western blotting.

### Lysate preparations

For immunoprecipitation, frozen RNAi worms were milled using the Mixer Mill MM 400 (Retsch; cycle at 25 times/sec for 2 min, 3 cycles total). Worm powder was thawed on ice and sonicated using an ultrasonic disruptor (UD201; Tomy Tech., Palo Alto, California, USA) with gauge at number 10, 5 times (20 sec followed by 1 min rest each time). To the lysate, 5 mM MgCl2 and 125 U/ml Benzonase were added, and the lysis was continued by rotating at 4 °C for 30 min. The lysate was then spun down at 22000 g for 20 min at 4 °C, and the supernatant was filtered with 40 μm cell strainer (Corning Falcon #352340, New York, USA) and saved at -80 °C.

To prepare samples for general western blotting of GFP-fused DSB-1, frozen worm pellet was suspended in urea lysis buffer (20 mM HEPES pH 8.0, 9M Urea, 1 mM sodium orthovanadate), sonicated (Taitec VP505 homogenizer, Koshigaya City, Japan; 50% output power, cycle of 10 sec on and 10 sec off for 7 min total) and spun down at 16000 g at 4 °C for 15 min. The supernatant was used to measure protein concentration using the BCA kit (Pierce BCA protein assay kit #23225; Thermo Scientific, Waltham, Massachusetts, USA), and a set amount of protein was loaded for western blotting after boiling for 10 min in SDS-PAGE sample buffer.

For endogenous DSB-1 immunoblotting experiments, 50 worms of 24 h post-L4 stage were picked into M9 + 0.05% Tween for each lane and washed twice, then SDS-PAGE sample buffer was added to the harvested worms, and after boiling for 5 min at 95°, the protein was flash centrifuged and loaded on the gel.

### Western blotting

For western blotting of GFP-fused DSB-1, SDS-PAGE was carried out using 5-12% Wako gradient gel (Wako #199-15191, Tokyo, Japan), and proteins were transferred to a PVDF membrane at 4 °C, 80 V for 2.5 h. The membrane was blocked with TBST buffer (TBS and 0.1% Tween) containing 5% skim milk (Nacalai Inc. #31149-75, Kyoto, Japan) at room temperature for 1 h and probed with primary antibody solution containing 2.5% skim milk at 4 °C overnight followed by additional 2 h at room temperature, washed with TBST for four times, probed with secondary antibody solution containing 2.5% skim milk at room temperature for 2 h, washed with TBST for four times. Chemi Luminol assay kit, Chemilumi-one super (Nacalai Inc. #02230-30, Kyoto, Japan) or Chemilumi-one ultra (Nacalai Inc. #11644-24, Kyoto, Japan), was used to visualize protein bands using an ImageQuant LAS4000 imager (GE Healthcare #28955810, Chicago, Illinois, USA).

For endogenous DSB-1 immunoblotting experiments, gel electrophoresis was performed using 4-12% Novex NuPage gels (Invitrogen #NP0322BOX, Waltham, Massachusetts, USA. Proteins were transferred to a PVDF membrane at 4 °C, 80 V for 2.5 h. The membrane was blocked at room temperature for 1 h in TBST containing 5% skim milk and then probed with primary antibody solution containing 5% skim milk at 4 °C overnight or at room temperature for 2 h, washed three times with TBST containing 1% skim milk, probed with secondary antibody solution containing 1% skim milk at room temperature for 2 h and washed with TBST for three times before proceeding to detection.

### Immunofluorescence and imaging

Immunostaining was performed as described in (70) with modifications as follows: Young adult worms (24 h post-L4 larval stage) were dissected in 15 μL EBT (27.5 mM HEPES pH 7.4, 129.8 mM NaCl, 52.8 mM KCl, 2.2 mM EDTA, 0.55 mM EGTA, 1% Tween, 0.15% Tricane) buffer, fixed by adding another 15 μL fixative solution (25 mM HEPES pH 7.4, 118 mM NaCl, 48 mM KCl, 2 mM EDTA, 0.5 mM EGTA, 1% Formaldehyde) and mixing for no more than 2 min in total on each coverslip. The excess liquid was pipetted off with 15 μL remaining which was picked up by touching a micro slide glass (Matsunami #S9901, Osaka, Japan) to the top of it before freezing at -80 °C. The slides were fixed in -20 °C methanol for exactly 1 min, transferred to PBST (PBS and 0.1% Tween) immediately and washed 3 times (10 min/time) by moving slides to fresh PBST at room temperature. Then the slides were blocked in 0.5% BSA in PBST for 30 min. Primary antibody incubation was performed at 4 °C overnight while secondary antibody incubation was performed for 2 h at room temperature. At last each slide was mounted with 15 μL mounting medium (250 mM TRIS, 1.8% NPG-glycerol) onto clean Matsunami No. 1 ½ (22 mm^2^) coverslip.

Images were captured by a Deltavision personalDV microscope (Applied Precision/GE Healthcare, Chicago, Illinois, USA) equipped with a CoolSNAP ES2 camera (photometrics) at a room temperature of 20–22°C, using a 100x UPlanSApo 1.4NA oil immersion objective (Olympus, Tokyo, Japan) and immersion oil (LaserLiquid; Cargille, Cedar Grove, New Jersey, USA) at a refractive index of 1.513. The Z spacing was 0.2 μm and raw images were subjected to constrained iterative deconvolution followed by chromatic correction. Image acquisition and deconvolution was performed with the softWoRx suite (Applied Precision/GE Healthcare, Chicago, Illinois, USA). Image postprocessing for publication was limited to linear intensity scaling and maximum-intensity projection using OMERO (71).

### Fluorescence in situ hybridization and quantification

The pairing on the right arm of chromosome V was monitored with fluorescence in situ hybridization (FISH) probes that label the 5S rDNA locus as described in (70) with modifications as follows: young adult worms (24 h post-L4 larval stage) were dissected in 15 μL EBT buffer and fixed by adding another 15 μL 1% paraformaldehyde for 1-2 min. The excess liquid was removed before freezing. The slides were fixed in -20 °C methanol for exactly 1 min, transferred to 2x SSCT (150 mM NaCl, 15 mM Na citrate pH 7, 0.1% Tween) immediately and washed 3 times (5 min/time) by moving slides to fresh 2x SSCT at room temperature. Next, the slides were put in a Coplin jar filled with EBFa (25 mM HEPES pH 7.4, 118 mM NaCl, 48 mM KCl, 2 mM EDTA, 0.5 mM EGTA, 3.7% formaldehyde) for another 5 min. After that, the slides were transferred to 2x SSCT and washed for 3 times (5 min/time) to remove the fixative. The slides were put into 50% formamide in 2x SSCT, incubated 10 min at 37 °C, and then transferred to a new jar with the same solution, incubated at 37 °C overnight. The probe solution (15 μL) was added onto a 22 × 22 mm coverslip. The worms on the slides were touched to the drop of probe solution on the coverslip until the liquid was spreaded out. After being sealed, the slides were denatured at 95 °C for 2 min. The slides were then washed with 2x SSCT, stained with DAPI, washed again with 2x SSCT, and mounted with 15 μL mounting medium onto clean Matsunami No. 1S (22 mm^2^) coverslip. Quantification of FISH foci was done as in (40). FISH probes were generated as previously described (2).

### RAD-51 foci quantification

Quantitative analysis of RAD-51 foci per nucleus was performed as in (40). For all the genotypes except for *rad-54* and *atm-1; rad-54* mutants, manual counting was performed. For *rad-54* mutants, semi-automated counting was used as below: for early zones (1 and 2) with very few RAD-51 foci, manual counting was performed. For zones 3 and above, programs (at github.com/pmcarlton/deltavisionquant) written in GNU Octave (72) were used to segment nuclei and count the number of RAD-51 foci in each nucleus. The programs proceed via the following steps: (1) nuclear centers are found by identifying all voxels above a threshold (calculated with the Otsu method (73) on a maximum-intensity projection image) that are also local maxima; these pixels are then subjected to a gravitational-type attraction that collapses clouds of pixels into small clusters that with few exceptions lie at the center of imaged nuclei. (2) The original positions of all the pixels that contributed to a cluster that fall within a given radius of the center are used to define a three-dimensional convex hull that represents the nuclear volume. (3) Positions of all RAD-51 foci are calculated by thresholding as in step 1. (4) RAD-51 foci are assigned to the convex hull in which they are enclosed. Detected foci not located inside any convex hulls are rejected as background spots. The convex hull outlines and number of foci per nucleus are displayed as 2D projections for each image data file, and used during visual inspection of the 3D data to correct or reject mistaken counts. The nuclei on the coverslip-proximal side of the gonads were scored for each genotype. Statistical comparisons were performed via two-tailed t test.

### DAPI body counting at diakinesis

For DAPI body counting, completely resolvable contiguous DAPI positive bodies were counted in three-dimensional stacks as described previously (40). With this criterion, chromosomes that happen to be touching can occasionally be counted as a single DAPI body.

### γ-irradiation

For DAPI body staining, late L4 larval stage worms were exposed to γ-rays for 58 minutes 30 seconds at 0.855 Gy/min (total exposure 50 Gy) in a Cs-137 Gammacell 40 Exactor (MDS Nordion, Ottawa, Canada). Irradiated worms were fixed 18-22 h after irradiation for DAPI staining, and imaged to score DAPI-stained bodies as above. For immunoprecipitation of GFP-fused DSB-1, 24 h post-L4 worms were exposed to γ-rays for 11 minutes 42 seconds (total exposure 10 Gy). For general western blotting of DSB-1, approximately 24 h post-L4 worms were irradiated with either 10 Gy or 100 Gy of γ-rays, and animals were lysed 1 h post irradiation.

### Embryonic viability scoring

To score embryonic viability and male progeny of each genotype, L4 larval stage hermaphrodites (P0s) were picked individually onto plates and transferred to fresh plates every 24 h for 5 days. Unhatched eggs remaining on the plates 20 h after being laid were counted as dead eggs every day. Viable F1 progeny and males were scored 4 days after P0s were removed from corresponding plates.

### Multiple sequence alignment

Protein sequences in the DSB-1/2 orthology group were retrieved from the Caenorhabditis Genomes Project (caenorhabditis.org). Due to the high diversity within this group, the list was pared down to DSB-1 orthologs of 11 species in the Elegans group (**Supplemental Figure 3**) with orthologs of DSB-2 and other proteins omitted. The protein prediction of DSB-1 for *C. latens* was found to be incomplete, so it was reconstructed by hand from the transcripts in Bioproject PRJNA248912, from WormBase ParaSite version 14 (74). The sequences were then aligned using the L-INS-i setting of mafft v7.487 (75).

### Orthology clustering and gene tree inference

We downloaded protein FASTA and GFF3 files for 39 Caenorhabditis species and two outgroup species (*Diploscapter coronatus* and *Diploscapter pachys*) from WormBase ParaSite (74) and the Caenorhabditis Genomes Project website (caenorhabditis.org). We used AGAT v0.4.0 (76) to select the longest isoform for each protein-coding gene, and clustered the filtered proteins into putatively orthologous groups (OGs) using OrthoFinder v2.5.4 (77), using an inflation value of 1.3. We identified the group containing the *C. elegans* DSB-1 and DSB-2 proteins, aligned the sequences using FSA v1.15.9 (78), and inferred a gene tree using IQ-TREE v2.2.0-beta (79) under the LG substitution model with gamma distributed rate variation among sites. We visualised the resulting gene tree using iTOL (80) and extracted branch lengths using the ETE3 Python module (81).

## Supporting information

resource-table

## ACKNOWLEDGEMENTS

We would like to thank Minami Murai, Takaya Hashimoto, Tjebbe Boersma, Yuji Yamauchi, Hao Li, and Yoko Fujita for technical assistance. Many nematode strains were provided by the Caenorhabditis Genetics Center, which is funded by the National Institutes of Health National Center for Research Resources.

---

**Supplemental Figure 1.**
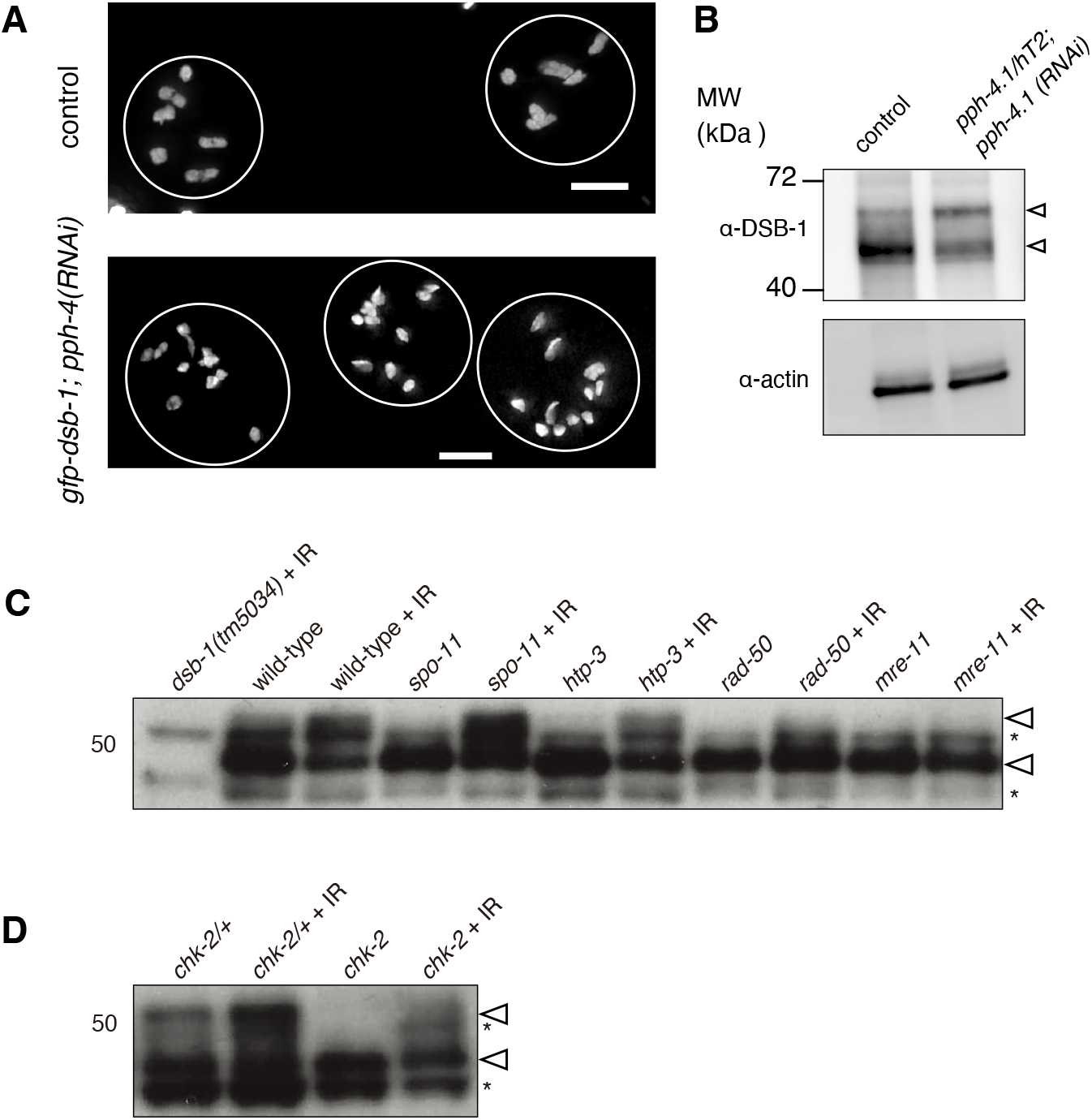
Validation of effectiveness of *pph-4* RNAi and western blots of DSB-1. **(A)** Images of DAPI-stained diakinesis nuclei of control (N2) or *gfp-dsb-1* treated with *pph-4* RNAi. The control nuclei contained 6 bivalents while the majority of nuclei contained univalents in *gfp-dsb-1; pph-4 (RNAi)* animals. **(B)** Western blot of endogenous DSB-1. Extracts of young adults (24 h post-L4 stage), from wild-type controls (left) or a mixed population of *pph-4*.*1(tm1598)* balanced heterozygous and homozygous mutants further treated with *pph-4*.*1* RNAi (right), probed with α-DSB-1. A total protein of 100 μg was loaded in each lane. Loading controls (α-actin) are shown at bottom. Arrowheads indicate two bands in the blot. **(C, D)** Western blots of endogenous DSB-1 in wild type (N2), *spo-11(me44), htp-3(tm3655), rad-50(ok197), mre-11(ok179)* or *chk-2(me64)* mutants in combination with 10Gy γ-irradiation as indicated. Lysate from 50 animals at 24 h post-L4 stage was loaded in each lane. Asterisks show two non-specific bands while arrowheads show two specific bands.

**Supplemental Figure 2.**
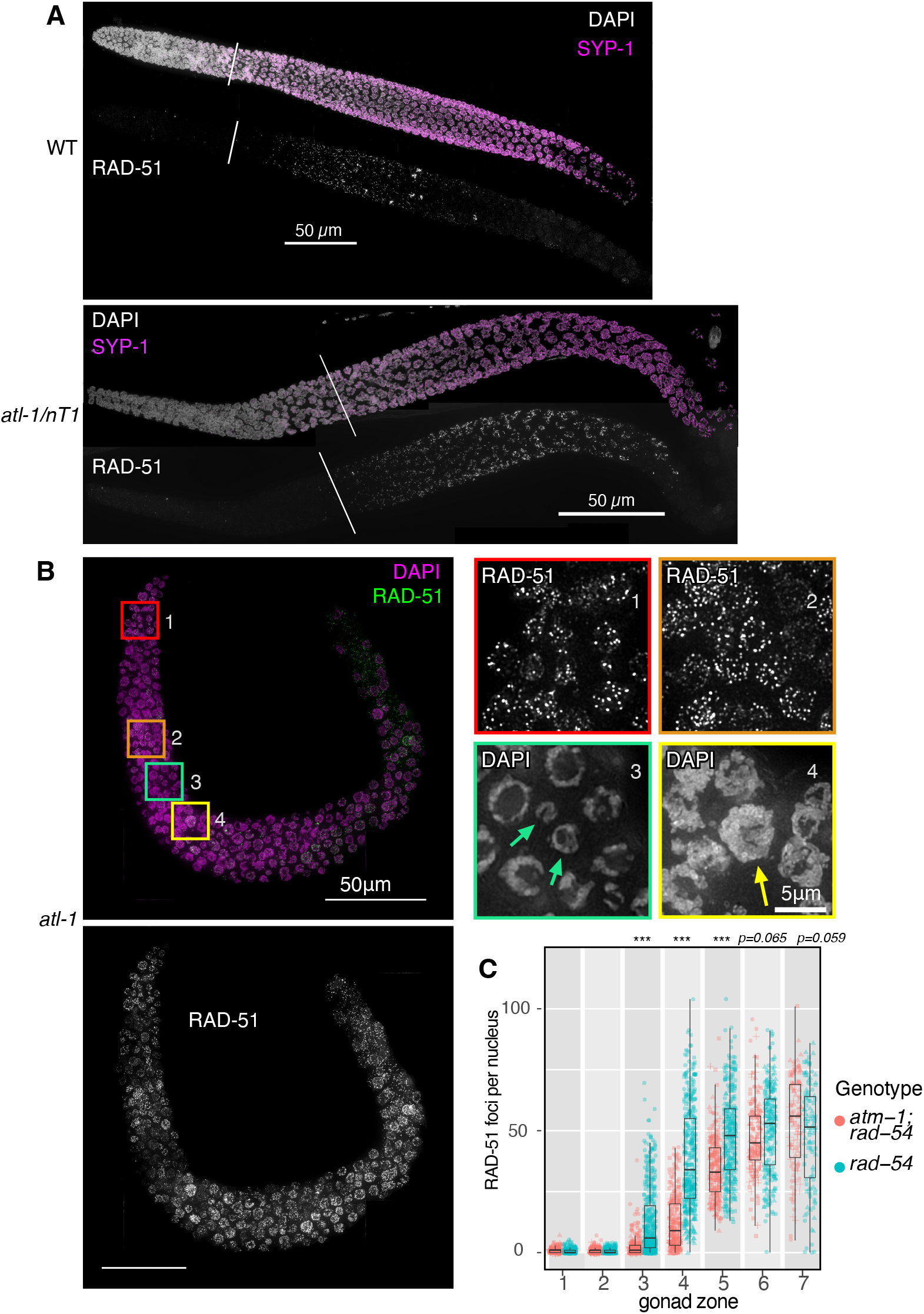
DSB formation in *atl-1* and *atm-1* mutants. **(A) *Top*:** Maximum-intensity projection of a wild type gonad. Above, DAPI staining is shown in grayscale and SYP-1, a component of the synaptonemal complex is shown in magenta. Below, RAD-51 is shown in grayscale. Diagonal line indicates the beginning of pachytene, before which few or no RAD-51 foci are seen. ***Bottom***: similar projection of a gonad from an *atl-1(tm853)/nT1* mutant. Scale bar, 50 μm. **(B)** Maximum-intensity projection of an entire gonad arm from an *atl-1(tm853)* mutant. Top left panel shows DAPI staining in magenta and RAD-51 staining in green. Bottom left panel shows RAD-51 foci in grayscale. Scale bar, 50 μm. Boxed insets on the top right show magnifications of the indicated color-matched regions highlighted on the left, which show numerous RAD-51 foci in both premeiotic and meiotic regions. Arrows indicate micronuclei resulting from improper mitotic division in box 3, and a large polyploid nucleus in box 4. Scale bar, 5 μm. **(C)** Quantitation of RAD-51 foci in each of seven zones, compared between *rad-54* single mutants and *atm-1(tm853); rad-54* double mutants. Significance was assessed via two-tailed *t* test, ***P<0.001.

**Supplemental Figure 3.**
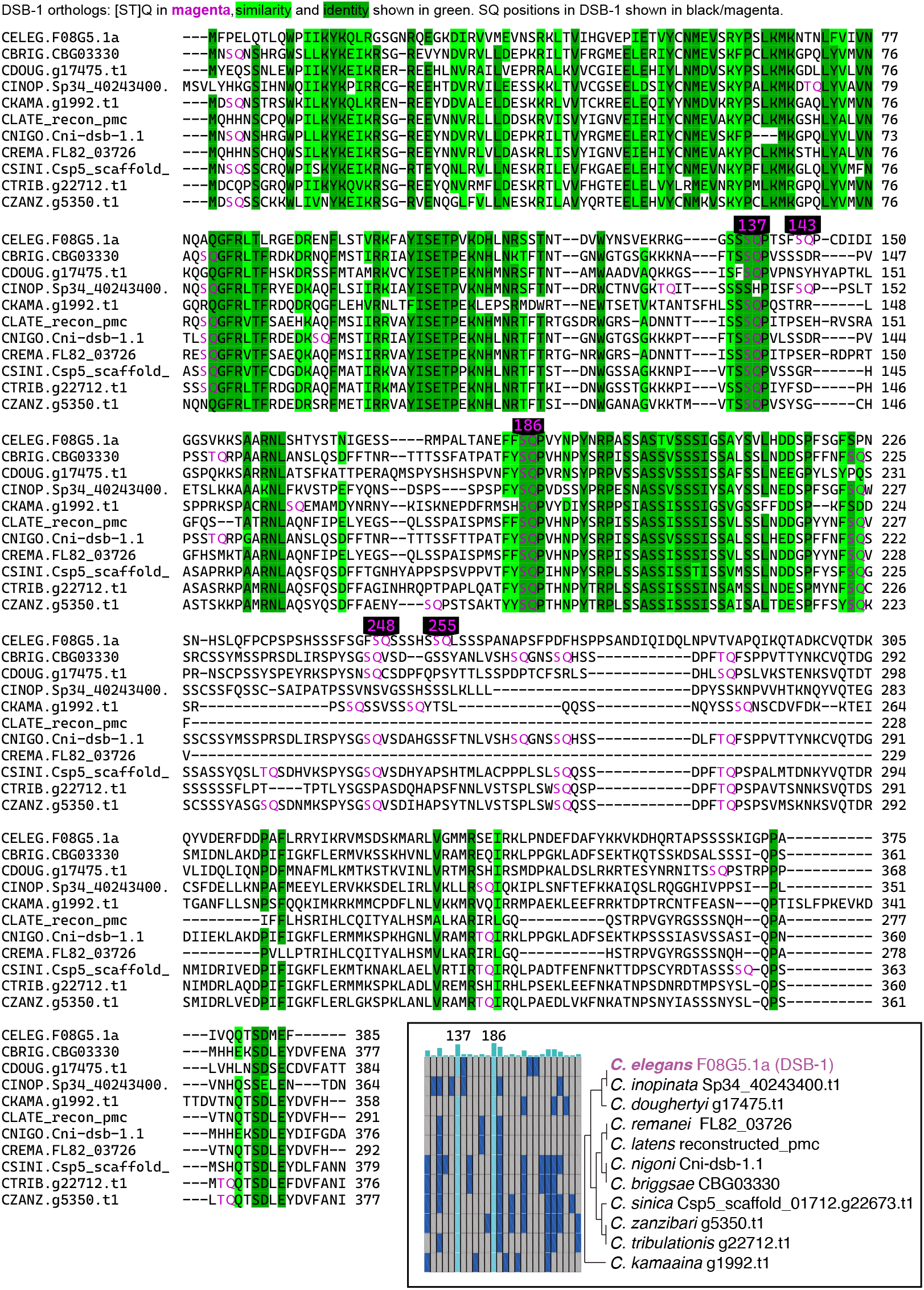
Sequence alignment of DSB-1 orthologs. Multiple sequence alignment (MSA) of DSB-1 orthologs in 11 Caenorhabditids. [ST]Q sites are depicted in magenta. Similarity and identity are shown in shades of green. The five SQ sites of DSB-1 in *C. elegans* are highlighted in black. ***Right bottom***: all [ST]Q sites in the 11 species shown as a grid: sorted by their position from left to right in the alignment, dark blue indicates presence of an [ST]Q site shared by up to 9 species; cyan indicates a site shared by 10 or 11 species. Species and protein names are shown at right, along with an unrooted, unscaled phylogenetic tree to show relatedness of each species (83).

**Supplemental Figure 4.**
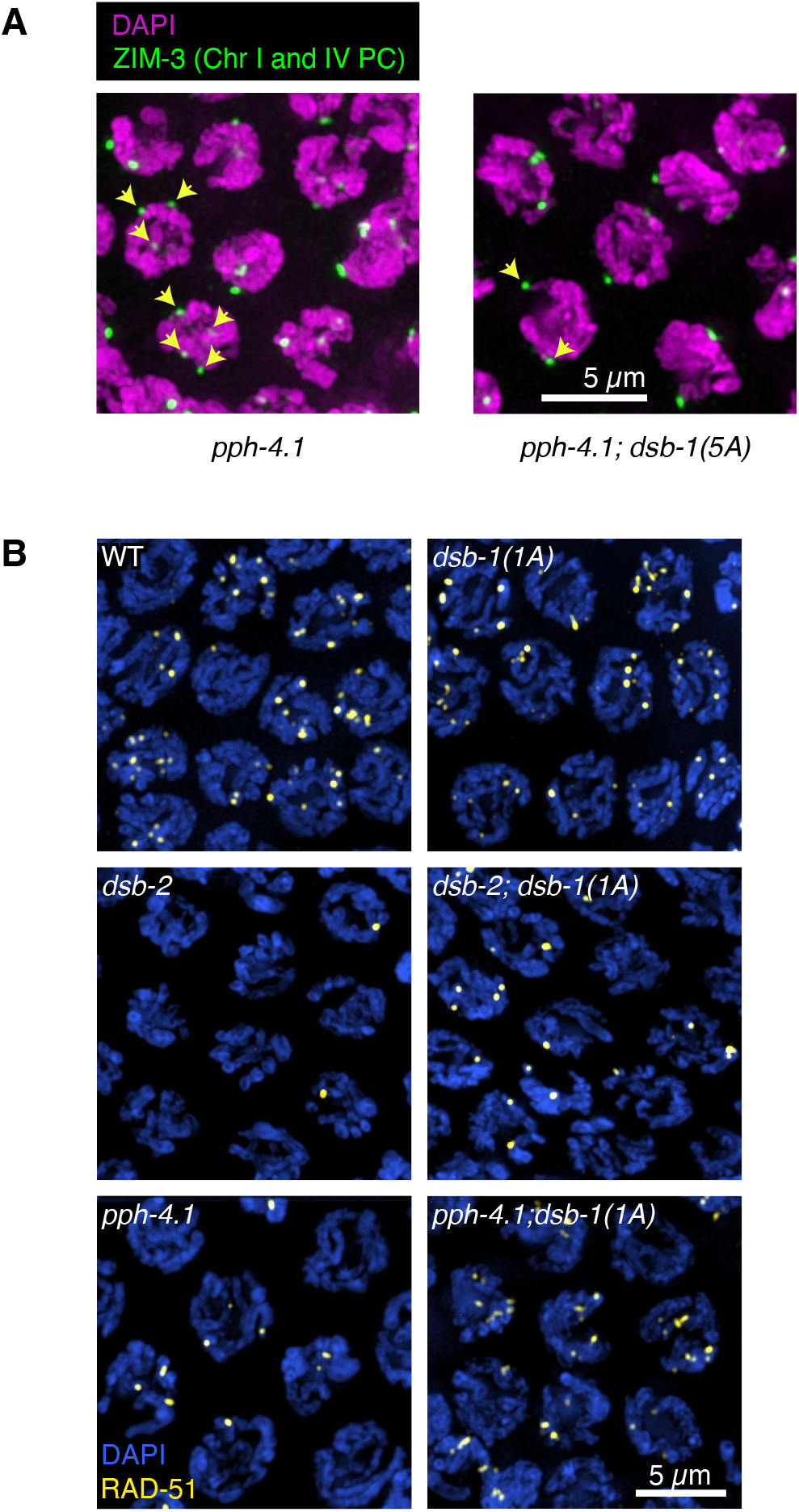
Homologous pairing in *pph-4*.*1; dsb-1(5A)* mutants and DSB formation in *pph-4*.*1; dsb-1(1A)* and *dsb-2; dsb-1(1A)* mutants. **(A)** Immunofluorescence image of ZIM-3 staining, detecting the pairing centers of chromosome I (right end) and IV (left end) in early pachytene in *pph-4*.*1(tm1598)* (left) and *pph-4*.*1(tm1598); dsb-1(5A)* (right) mutants. Maximum-intensity projections of DAPI are shown in magenta; ZIM-3 staining is shown in green. Yellow arrows point to examples of 3 or 4 foci per nucleus in *pph-4*.*1(tm1598)* single mutants (left); double mutants of *pph-4*.*1(tm1598)*; *dsb-1(5A)* show increased number of paired ZIM-3 foci (2 foci per nucleus), indicating rescued homologous pairing of chromosomes I and IV. Scale bar, 5 μm. **(B)** Immunofluorescence images of wild type, *dsb-1(1A), dsb-2(me96), dsb-2(me96); dsb-1(1A), pph-4*.*1(tm1598)*, and *pph-4*.*1(tm1598); dsb-1(1A)* mutants. Maximum-intensity projections of nuclei at mid-pachytene are shown with DAPI staining in blue and RAD-51 staining in yellow. Scale bar, 5 μm.

**Supplemental Figure 5.**
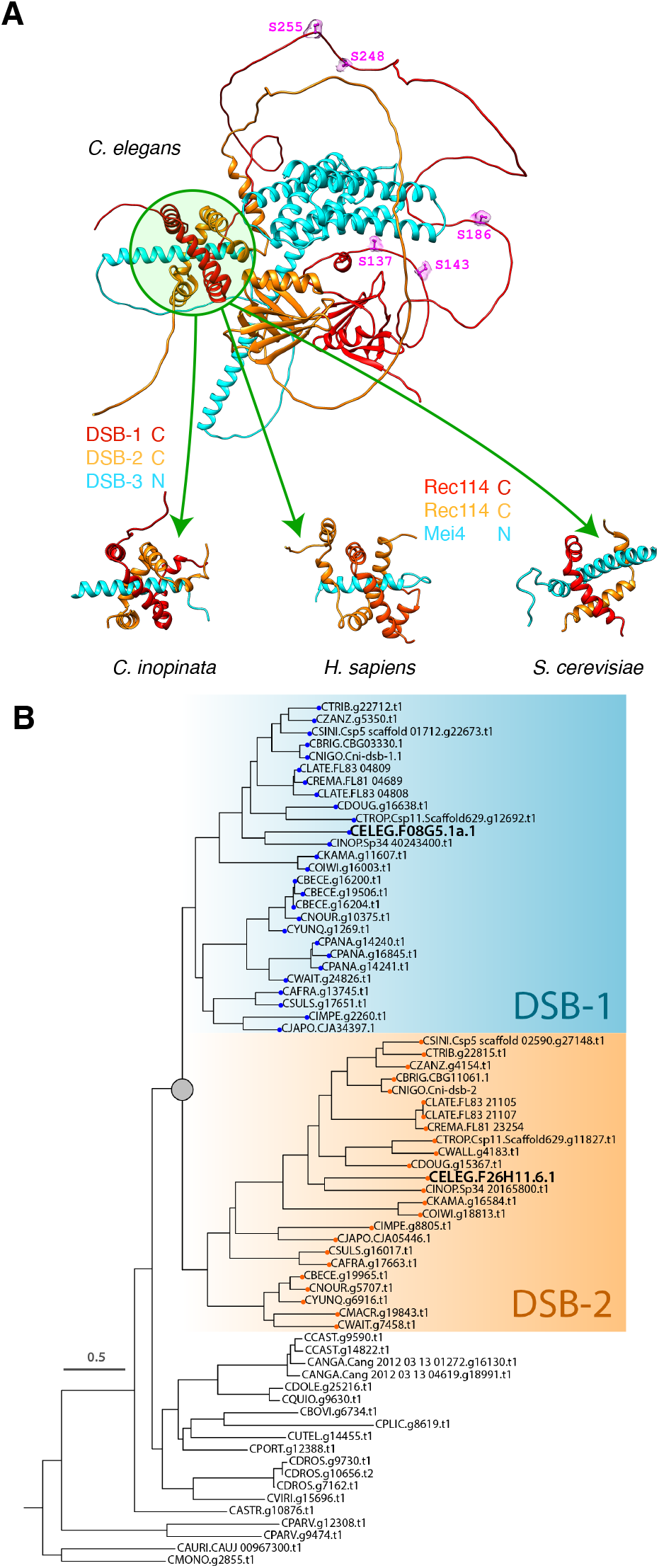
Structural and phylogenetic predictions of double-strand break factors. **(A)** A representative structure of the DSB-1, DSB-2, and DSB-3 heterotrimer predicted by the AlphaFold structure prediction pipeline (61, 62) is shown at top. Green circle highlights the region predicted to be the trimerization interface containing the C-termini of DSB-1 and -2, and the N-terminus of DSB-3. ATM/ATR kinase phosphorylation consensus sites in the predicted disordered loop of DSB-1 are shown in magenta and labeled. Sub-regions of similar structures predicted for orthologs of DSB-1, DSB-2, and DSB-3 in the *C. elegans* sister species *Caenorhabditis inopinata*, as well as the putative Rec114/Rec114/Mei4 heterotrimer in human and budding yeast, are shown below. In all cases a predicted N-terminal alpha-helix of the Mei4 ortholog (DSB-3) transfixes a channel formed by the predicted C-terminal helices of the Rec114 orthologs wrapping around each other. **(B)** Maximum likelihood gene tree of DSB-1/2 homologs in the genus Caenorhabditis. Gene tree of the orthogroup containing the *C. elegans* proteins DSB-1 (CELEG.F08G5.1a) and DSB-2 (CELEG.F26H11.6) was inferred using maximum likelihood (LG substitution model with gamma distributed rate variation). The DSB-1/DSB-2 duplication event is denoted by a gray circle. Branch lengths represent the number of substitutions per site; scale is shown at left. The tree is rooted on the branch subtending the *C. monodelphis* and *C. auriculariae* sequences.

